# Sphingolipid-Induced Programmed Cell Death Is a Salicylic Acid and EDS1-Dependent Phenotype in Arabidopsis

**DOI:** 10.1101/2021.04.20.440624

**Authors:** Stefanie König, Jasmin Gömann, Agnieszka Zienkiewicz, Krzysztof Zienkiewicz, Dorothea Meldau, Cornelia Herrfurth, Ivo Feussner

## Abstract

Ceramides and long chain bases (LCBs) are plant sphingolipids involved in the induction of plant programmed cell death (PCD). The *fatty acid hydroxylase* mutant *fah1 fah2* exhibits high ceramide levels and moderately elevated LCB levels. Salicylic acid (SA) is strongly induced in these mutants, but no cell death is visible. To determine the effect of ceramides with different chain lengths, *fah1 fah2* was crossed with *ceramide synthase* mutants *longevity assurance gene one homologue1-3* (*loh1*, *loh2* and *loh3*). Surprisingly, only triple mutants with *loh2* show a cell death phenotype under the selected conditions. Sphingolipid profiling revealed that the greatest differences between the triple mutant plants are in the LCB and LCB-phosphate (LCB-P) fraction. *fah1 fah2 loh2* plants accumulate LCB d18:0 and LCB-P d18:0. Crossing *fah1 fah2 loh2* with the SA synthesis mutant *sid2-2*, and with the SA signaling mutants *enhanced disease susceptibility 1-2* (*eds1-2*) and *phytoalexin deficient 4-1* (*pad4-1*), revealed that lesions are SA- and EDS1-dependent. These quadruple mutants also suggest that there may be a feedback loop between SA and sphingolipid metabolism as they accumulated less ceramides and LCBs. In conclusion, PCD in *fah1 fah2 loh2* is a SA and EDS1-dependent phenotype, which is likely due to accumulation of LCB d18:0.

## Introduction

Sphingolipids are important molecules with diverse functions in eukaryotic cells. In addition to their structural role as membrane components, in plants they are critical for development and responses to biotic and abiotic stresses (Ali et al. 2018; Berkey et al. 2012; Huby et al. 2020; Saucedo-Garcia et al. 2015). Plant sphingolipids can be divided into four subgroups: free long chain bases (LCBs), ceramides (Cers), glucosylceramides (GlucCers) and glycosyl inositol phosphoryl ceramides (GIPCs) (Pata et al. 2010). Whereas GlucCers and GIPCs are important structural membrane components, Cers and LCBs are mainly described as biosynthetic intermediates and signaling molecules.

LCBs consist of a C18 core with an amino group at C2. Hydroxylation at C1 and C3 generates sphinganine (d18:0), the simplest form of LCB. It can be directly channeled into ceramide synthesis, or modified by additional hydroxylation at C4 (t18:0) or desaturation (d18:1) (Luttgeharm et al. 2016b). Ceramide Synthase (CerS) catalyzes the step from LCB to ceramide by *N*-acylation of the LCB (Luttgeharm et al. 2016b; Michaelson et al. 2016). The fatty acid moiety can vary in chain length between C16 and C26. In Arabidopsis there are three different CerS enzymes: Longevity Assurance Gene One (LAG1) Homologue1-3 (LOH1-3) (Markham et al. 2011; Ternes et al. 2011). LOH1 and LOH3 preferentially catalyze the reaction with very long chain fatty acids (VLCFAs) and t18:0 LCBs as substrates (Luttgeharm et al. 2016a; Markham et al. 2011; Ternes et al. 2011). LOH2 catalyzes the reaction preferentially with long chain fatty acids (LCFAs) of C16 chain length and d18:0 LCB (Luttgeharm et al. 2016a; Markham et al. 2011; Ternes et al. 2011). Consequently, *loh1* mutants accumulate Cers and GlucCers with C16 fatty acid and free trihydroxy sphingoid bases, and show lesions late in development, whereas *loh2* mutants show reduced LCFA-containing Cer levels but exhibit no apparent growth phenotype (Markham et al. 2011; Ternes et al. 2011). These data suggest that spontaneous cell death in *loh1* mutants is triggered either by the accumulation of free trihydroxy LCBs or C16 ceramide species (Ternes et al. 2011).

The fatty acid moiety of a Cer molecule can be modified by hydroxylation in the alpha position. In Arabidopsis, two Fatty Acid Hydroxylases (FAH) are described (FAH1 and FAH2) (König et al. 2012; Nagano et al. 2009). FAH1 prefers ceramides with VLCFAs and FAH2 with C16 fatty acids (Nagano et al. 2012). Hydroxylated Cers (hCers) are found in GlucCers and GIPCs in high amounts (Pata et al. 2010) and might be suppressors of cell death (Townley et al. 2005). In *fah1 fah2* double mutant plants, however, high amounts of non-hydroxylated Cers in relation to hCers were found but no cell death was observed (König et al. 2012). Additionally, free SA and SA glucoside (SAG) accumulate in those plants.

Apart from *loh* and *fah* mutants, deregulation of sphingolipid synthesis in other steps of the pathway often leads to cell death with involvement of SA (Bi et al. 2014; Li et al. 2016; Yanagawa et al. 2017; Zienkiewicz et al. 2020). Different Arabidopsis mutants with defects in sphingolipid metabolic pathways spontaneously accumulate SA and undergo programmed cell death (PCD) (Bi et al. 2014; Brodersen et al. 2002; Greenberg et al. 2000; Wang et al. 2008; Zienkiewicz et al. 2020). So far, the regulation of SA accumulation by sphingolipids on the molecular level is mostly unknown, but there are many hints that Cers and /or LCBs are inducers of PCD signaling and SA induction.

In this study, we examined why *fah1 fah2* mutants do not display cell death, and which components of the sphingolipid pool might be responsible for PCD. We further looked into the involvement of SA in sphingolipid-induced cell death by crossing sphingolipid-deficient mutants with SA biosynthesis or signaling mutants.

## Results

### *Crossing* loh2 *into* fah1 fah2 *mutants leads to lesions in distinct leaf areas*

The *fah1 fah2* double mutant has elevated Cer levels and slightly elevated LCB levels, but no severe lesion phenotype (König et al. 2012). To further analyze if Cers with distinct chain lengths may lead to lesions in these mutants, as previous work suggested (Ternes et al. 2011), the double mutant was crossed with the three CerS mutants *loh1*, *loh2* and *loh3*.

Crossing *fah1 fah2* with *loh1* and *loh3* produced triple mutants with no additional severe phenotypes. However, crossing *loh2* into *fah1 fah2* produced triple mutants with lesions at distinct areas at the leaves (Fig. S1B), which was verified by trypan blue staining (Fig. 1). Additionally, these plants are significantly smaller than the double mutant plants. Whereas *fah1 fah2* plants are more than 50% reduced in rosette area compared to Col-0 mutant plants, *fah1 fah2 loh2* plants show less than 30% of the rosette leaf area of wild type plants (Fig. S1A). *fah1 fah2 loh1* and *fah1 fah2 loh3* mutants do not show visible lesions or changes in leaf area compared to *fah1 fah2* grown for up to 35 days under long day conditions.

**Fig. 1.**
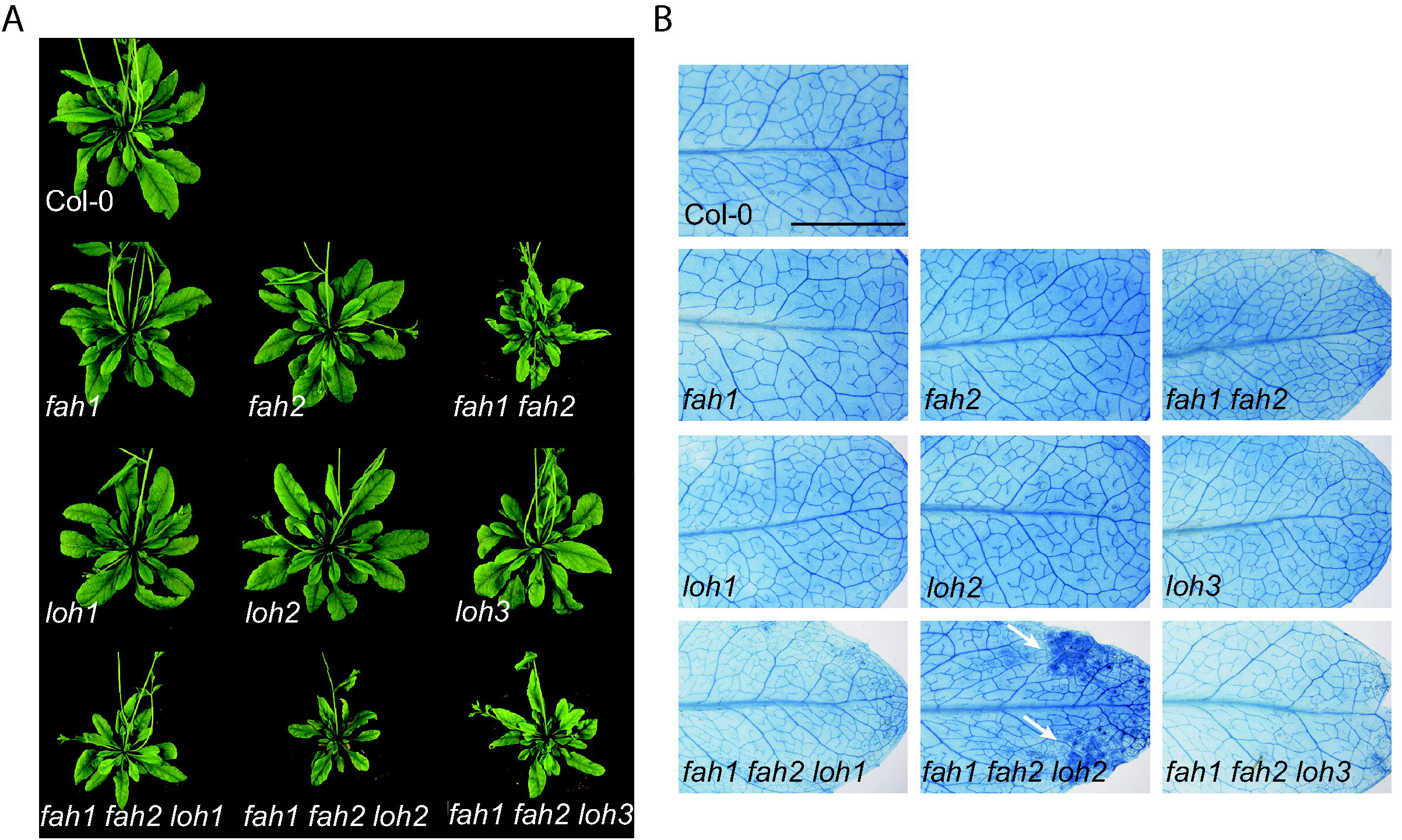
Crosses of *fah1 fah2* double mutants with CerS mutants *loh1 - 3* leads to a lesion phenotype in *fah1 fah2 loh2* plants. (A) Representative images of single, double and triple mutant plants grown for 35 days under long day conditions. (B) Trypan blue staining of single leaves of 27-day-old plants grown under long day conditions (bar, 5 mm). White arrows show lesions in *fah1 fah2 loh2* plants. Representative leaves out of 5 replicates are shown.

### fah1 fah2 loh2 *triple mutants accumulate d18:0 and d18:0-P*

To reveal the reason for lesion development in *fah1 fah2 loh2* mutant plants, sphingolipid profiles in all mutant lines were compared. Before analyzing large data sets, Cer and SA levels at early (14-day-old) and late time points (35-day-old) were compared to establish the best plant age for measurements of these metabolites (Fig. S2). SA levels between Col-0, *fah1 fah2*, *loh2* and *fah1 fah2 loh2* are similar at both time points, but highest at 35 days. However, changes in Cer and hCer profiles are more distinct in 35-day-old plants. Thus, plants at this age were used for further analysis.

Analysis of all mutant lines revealed that the most obvious changes between the triple mutants are in their LCB profiles. As shown in earlier studies, changes in LCB levels in the *fah1 fah2* double mutant are only minor (1.5-2-fold increase) compared to Col-0. However, *fah1 fah2 loh2* triple mutants accumulate d18:0 (20-fold compared to Col-0) and d18:0-P (not detectable in Col-0) as well as t18:0 (7-fold compared to Col-0) (Fig. 2A). In contrast, in *fah1 fah2 loh1* triple mutants t18:0 (7-fold compared to Col-0) and t18:1 (4.7-fold) are the dominant enriched species, but the amount is comparable to the *loh1* single mutant. *fah1 fah2 loh3* triple mutants do not show significant changes compared to *fah1 fah2* mutant LCB levels.

**Fig. 2.**
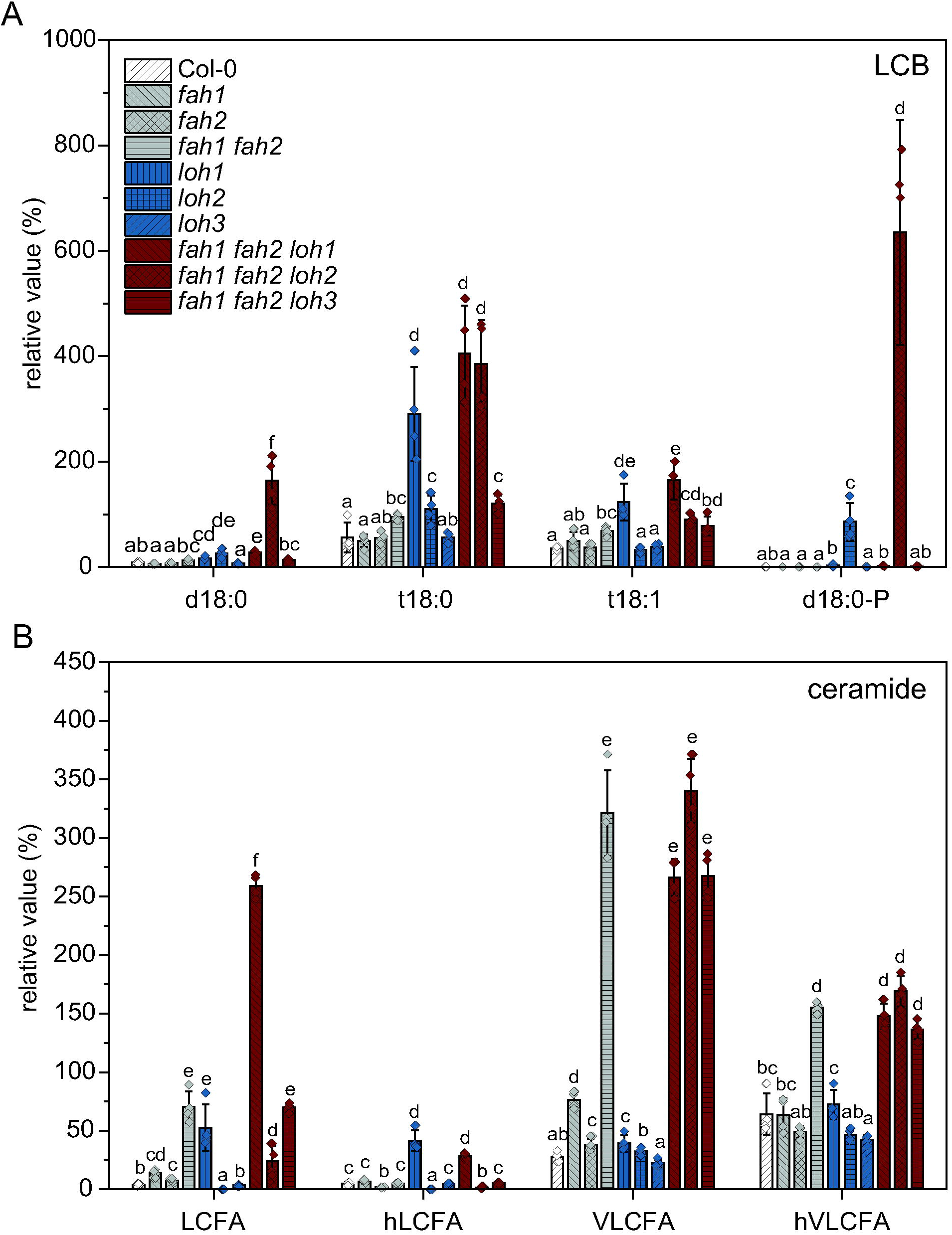
Triple mutant plants defective in CerS *LOH1* or *LOH2* and both *Fatty Acid Hydroxylases* (*fah1*, *fah2*) show altered LCB (A) and Cer (B) levels compared to wild type (Col-0) and to *Fatty Acid Hydroxylase* double mutant (*fah1 fah2*). Rosette leaves of 35-day-old plants grown under long day conditions were extracted and analyzed. LCFA, Cers with C16-C18 FA moiety; hLCFA, Cers with C16-C18 α-hydroxylated FA moiety; VLCFA, Cers with C20-C28 FA moiety; hVLCFA, Cers with C20-C28 α-hydroxylated FA moiety. Values represent the mean ±SD of four biological replicates (n=4). The experiment was repeated once with similar tendencies. Statistical analysis was performed with log-transformed data by one-way analysis of variance (ANOVA) with Tukey’s *post hoc* test (*P*<0.05). Different letters indicate significant differences with *P*<0.05.

In addition to LCBs, Cers are suspected to be involved in cell death induction in leaves. Changes in Cer profiles are especially visible in the *fah1 fah2 loh1* triple mutant (Fig. 2B). Compared to the *fah1 fah2* mutant, LCFA (C16-C18)-containing Cers increased up to three times, whereas in the *fah1 fah2 loh2* triple mutant these species are strongly reduced. VLCFA-containing Cers are comparable in the double and triple mutants.

### *SAG levels are highly elevated in mutants with* fah1 fah2 *background*

As SA is described to be involved in induction of sphingolipid derived cell death, SA and SAG were measured in rosette leaves. Among all mutant lines tested, the highest SA levels are detectable in the *fah1 fah2 loh2* mutant (5.3 nmol/g FW) followed by the *loh1* single mutant with 3.8 nmol/g FW (Fig. 3A). Compared to 1.7 nmol/g FW in Col-0, the induction is minor. However, SAG strongly accumulates in the *fah1 fah2* double mutant (105 nmol/g FW) and in all three triple mutant lines (*fah1 fah2 loh1*: 124 nmol/g FW, *fah1 fah2 loh2*: 188 nmol/g FW, *fah1 fah2 loh3*: 205 nmol/g FW) compared to Col-0 (17 nmol/g FW; Fig. 3B).

**Fig. 3.**
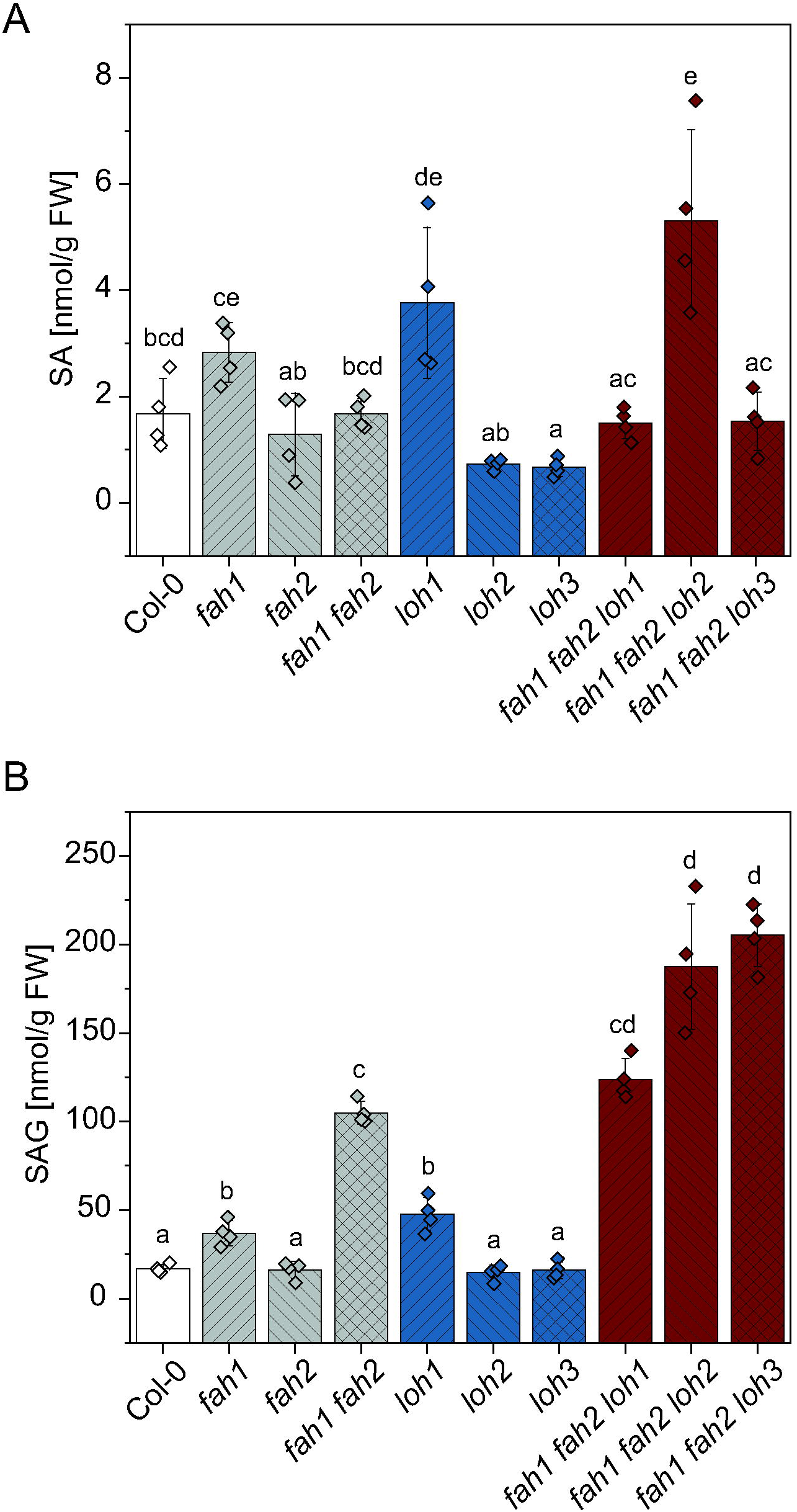
SA (A) and SAG (B) levels are altered in CerS (*loh1*, *loh2*, *loh3*) and *Fatty Acid Hydroxylase* (*fah1*, *fah2*) mutant plants and crosses. Rosette leaves of 35-day-old plants grown under long day conditions were extracted and analyzed. Values represent the mean ±SD of four biological replicates (n=4). The experiment was repeated once with similar tendencies. Statistical analysis was performed with log-transformed data by one-way analysis of variance (ANOVA) with Tukey’s *post hoc* test (*P*<0.05). Different letters indicate significant differences with *P*<0.05.

### *Reduction of SA levels by disruptions in SA synthesis or signaling represses the lesion phenotype of the* fah1 fah2 loh2 *mutant*

To connect SA accumulation with the cell death phenotypes, mutants were crossed with the SA biosynthesis mutant *sid2-2* (defective in *ISOCHORISMATE SYNTHASE 1*), and additionally with *ENHANCED DISEASE SUSCEPTIBILITY1-2* (*eds1-2*) and *PHYTOALEXIN DEFICIENT4* (*pad4-1*). *eds1-2* and *pad4-1* are signaling mutants defective in the induction of SA synthesis by effector-triggered immunity (ETI) (Cui et al. 2017).

After crossing the *fah1 fah2 loh2* triple mutant with *pad4-1* the lesion phenotype is still visible but less prominent than in *fah1 fah2 loh2* mutant plants (Fig. 4B). In contrast to this, in crossings of *fah1 fah2 loh2* triple mutant with *eds1-2* or *sid2-2* no lesions can be detected. Rosette leaf areas of all quadruple mutants are smaller than the respective single mutant (*fah1 fah2 loh2 pad4-1*: 58 %, *fah1 fah2 loh2 eds1-2*: 46 %, *fah1 fah2 loh2 sid2-2*: 34 %) but size reduction is not as strong as for the *fah1 fah2 loh2* mutant compared to Col-0 (73%; Fig. 4A, S1). Especially *fah1 fah2 loh2 sid2-2* plants show only a small reduction in size and resemble the phenotype of the *sid2-2* single mutant (Fig. 4A, S1).

**Fig. 4.**
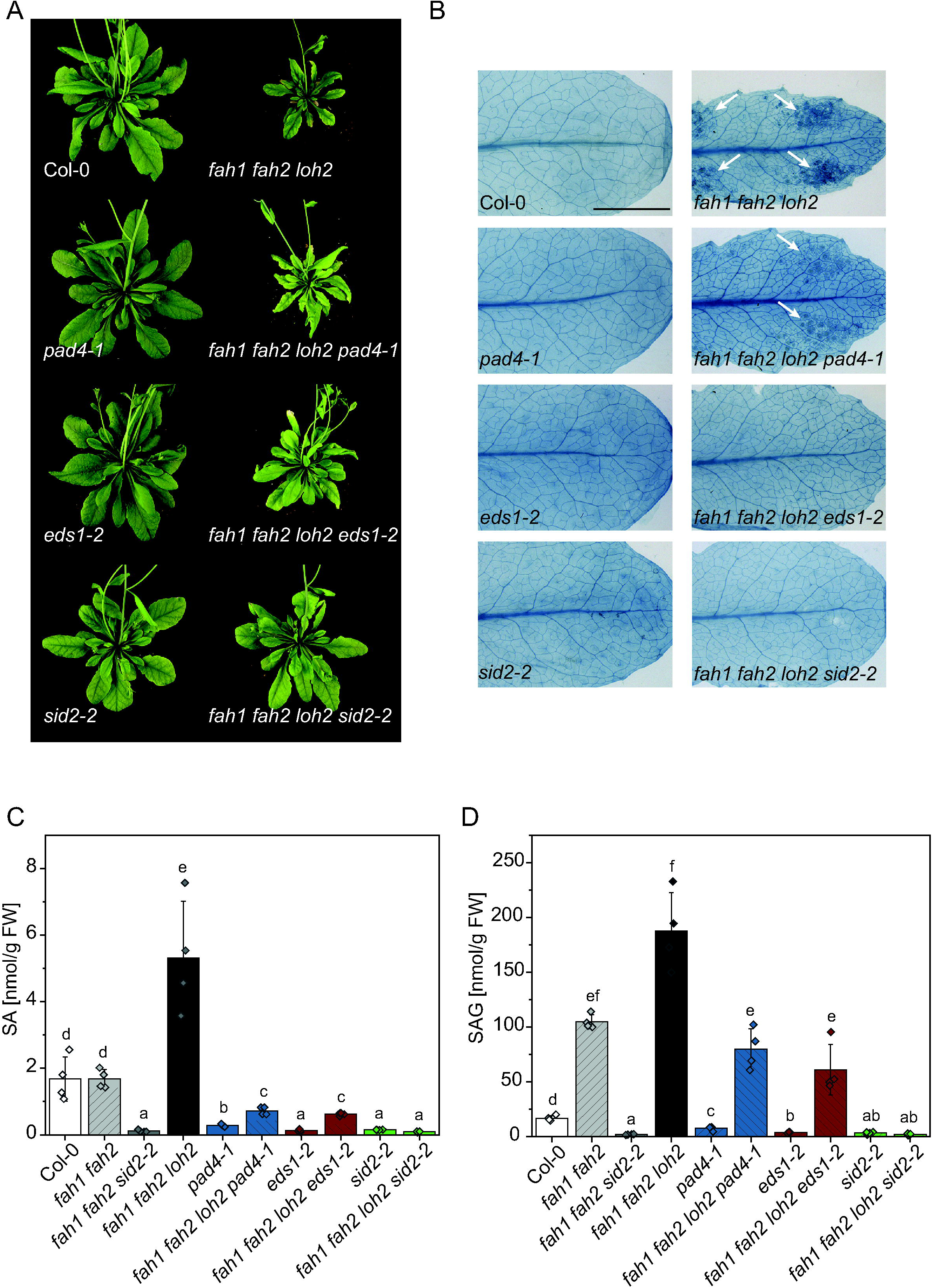
Crosses of *fah1 fah2 loh2* triple mutants with SA synthesis (*sid2-2*) and signaling mutants (*eds1-2, pad4-1*) show changes in lesion phenotype. (A) Representative images of single, triple and quadruple mutant plants grown for 35 days under long day conditions. (B) Trypan blue staining of single leaves of 25-day-old plants grown under long day conditions (bar, 5 mm). White arrows show lesions in *fah1 fah2 loh2* and *fah1 fah2 loh2 pad4-1* plants. Representative leaves out of four replicates are shown. The experiment was repeated twice with similar results. (C, D) SA and SAG levels of rosette leaves of 35-day-old plants. Mean values ±SD of four biological replicates (n=4) are shown. The experiment was repeated once with similar tendencies. Statistical analysis was performed with log-transformed data by one-way analysis of variance (ANOVA) with Tukey’s *post hoc* test (*P*<0.05). Different letters indicate significant differences with *P*<0.05.

To confirm the reduction of SA and SAG in the mutants, these phytohormones were measured in all mutant lines. SA levels in the *fah1 fah2 loh2* plants crossed with *eds1-2* and *pad4-1* are lower than wild type levels. SAG is still induced in those plants, but at a lower level than in *fah1 fah2 loh2* plants (*fah1 fah2 loh2 pad4-1*: 43% of *fah1 fah2 loh2* levels, *fah1 fah2 loh2 eds1-2*: 33% of *fah1 fah2 loh2* levels; Fig 4C). *fah1 fah2 loh2* plants crossed with *sid2-2* plants show no SA or SAG accumulation. In addition to the *fah1 fah2 loh2* triple mutant the *fah1 fah2* double mutant was also crossed with *sid2-2* to analyze which phenotypic changes are due to SA accumulation in those plants. Also in this mutant line, no SA or SAG is detectable.

Sphingolipid analyses of the mutants show that defects in SA synthesis and/or signaling also have an influence on the sphingolipidome of the plants. All SA mutant crosses showed a reduction of LCB accumulation relative to the *fah1 fah2 loh2* triple mutant. d18:0 and d18:0-P are especially strongly reduced in the quadruple mutants (Fig. 5A). Additionally, LCFA-containing Cers are depleted in the quadruple mutants, and reductions of VLCFA-containing Cer are only minor (Fig. 5B). In contrast to this, *fah1 fah2* mutants crossed with *sid2-2* show a strong reduction in VLCFA-containing Cers and only a slight reduction in LCB levels compared to *fah1 fah2* (Fig. 5B). In contrast to LCBs and Cers, levels of GlucCer and GIPC are not changed due to disruption of SA synthesis (Fig. S3).

**Fig. 5.**
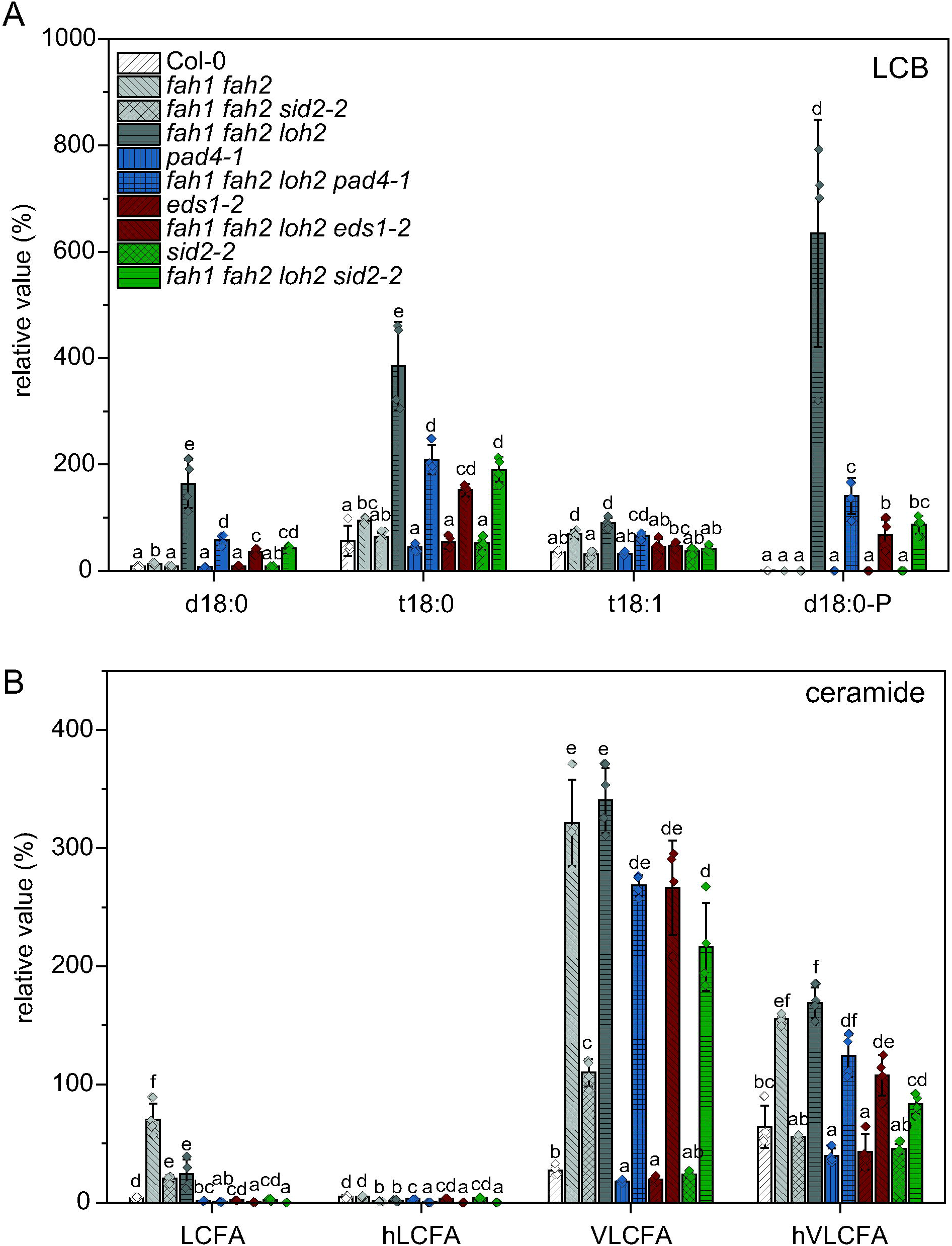
LCB (A) and Cer (B) profiles are altered in crosses of *fah1 fah2* double and *fah1 fah2 loh2* triple mutants with SA synthesis (*sid2-2*) and signaling mutants (*eds1-2*, *pad4-1*). Rosette leaves of 35-day-old plants grown under long day conditions were extracted and analyzed. LCFA, Cers with C16-C18 FA moiety; hLCFA, Cers with C16-C18-hydroxylated FA moiety; VLCFA, Cers with C20-C28 FA moiety; hVLCFA, Cers with C20-C28 α hydroxylated FA moiety. Values represent the mean ±SD of four biological replicates (n=4). The experiment was repeated once with similar tendencies. Statistical analysis was performed with log-transformed data by one-way analysis of variance (ANOVA) with Tukey’s *post hoc* test (*P*<0.05). Different letters indicate significant differences with *P*<0.05.

## Discussion

Accumulation of LCBs and/or Cers is suspected to be responsible for sphingolipid-induced cell death in several mutants with disruptions in the sphingolipid pathway (Berkey et al. 2012; Bi et al. 2014; Yanagawa et al. 2017; Zienkiewicz et al. 2020). However, *fah1 fah2* mutant plants exhibit strong accumulations of non-hydroxylated Cers, hVLCFA-containing Cers, moderate increases in LCB levels, and enhanced SA levels, but no lesion phenotype (König et al. 2012). Therefore, the intention behind crossing *fah1 fah2* plants with *loh* plants was to determine if specific changes in the sphingolipidome of those mutants could be correlated with the induction of PCD. *fah1 fah2* mutants were crossed with *loh1, loh2* and *loh3* mutants to shift the Cer accumulation to VLCFA Cer or C16 Cer. C16 Cers are discussed as PCD-inducing sphingolipids in *loh1* plants under certain conditions (Ternes et al. 2011) and in *LOH2*-overexpressing plants (Luttgeharm et al. 2015). Unexpectedly, *fah1 fah2* crosses with *loh2*, not *loh1*, produced triple mutants with PCD (Figs. 1 and 2). In *fah1 fah2 loh2* plants LCFA-containing Cers are strongly reduced compared to *fah1 fah2* but this is also more pronounced in the *loh2* single mutant. Therefore, it seems that the other main difference in the sphingolipid profile of these plants, the elevated levels of d18:0 LCB and d18:0-P LCB, are most likely the underlying reason for the lesion phenotype. LCBs are also suspected to be involved in PCD induction upon Fumonisin B_1_ treatment, a fungal toxin inhibiting LOH1 (Luttgeharm et al. 2016a). LOH1 inhibition induces a strong accumulation of d18:0 and t18:0 as well as LCFA-containing Cer species. This is supported by the analysis of mutants defective in different steps of the sphingolipid pathway or pathogen-infected plants, which collectively suggest that accumulation of LCBs induces cell death in plants (Luttgeharm et al. 2016a; Peer et al. 2010; Saucedo-Garcia et al. 2015; Shi et al. 2007; Yanagawa et al. 2017). The different roles of d18:0 and t18:0 LCBs are still not clear. In case of *fah1 fah2 loh2* both species are induced, but compared with *fah1 fah2 loh1* accumulation of d18:0 LCB is specific to the former. d18:0 LCB may therefore be responsible for the differences in lesion development between the mutants. It could also be that d18:0 LCB is acting together with t18:0 LCB and that the combined accumulation of both LCBs passes a critical level that leads to induction of PCD. LCB-Ps are thought to be the inactivated form of LCB that do not induce PCD and do not influence LCB-induced cell death (Glenz et al. 2019; Yanagawa et al. 2017). The high accumulation of d18:0-P in *fah1 fah2 loh2* plants might be due to the partial removal of d18:0 by phosphorylation by LCB kinase to avoid the harmful effect of the molecule (Imai and Nishiura 2005).

LOH2 seems not to be essential under normal growth conditions, as *loh2* mutants do not show a phenotype compared to wild type (Luttgeharm et al. 2015; Markham et al. 2011; Ternes et al. 2011). However, overexpression of *LOH2* leads to dwarfed plants with high amounts of LCFA-containing Cers and to higher resistance to FB_1_ treatment (Luttgeharm et al. 2015). Therefore, the authors suspected that LOH2 might be a safety valve for plants to sequester excess LCBs. In the case of the *fah1 fah2* double mutant LOH2 might be essential to channel excess LCBs into the Cer pool to avoid toxic LCB levels in the plant. In triple mutants with *loh1* or *loh3*, activity of LOH2 and the other remaining LOH are high enough not to exceed the critical level of LCBs. In the *fah1 fah2 loh2* mutant, this safety valve is not functional anymore as Cer synthesis of C16 fatty acids with d18-0 LCB is blocked and d18:0 LCB accumulate. Excess d18:0 can then be partially removed by phosphorylation resulting in accumulation of d18:0-P. It might be interesting to determine if blocking phosphorylation, for example in LCB kinase mutants, would lead to a more severe phenotype.

All mutants with *fah1 fah2* background showed high levels of total SA. To test the role of SA in the development of lesions and reduction in rosette size, mutants were crossed with the SA synthesis mutant *sid2-2* (isochorismate synthase 1 deficient). Additionally, as it is known that sphingolipid metabolism in plants is connected to plant defense (Huby et al. 2020), plants were crossed with *eds1-2* and *pad4-1*. EDS1 and PAD4 are necessary for promotion of *ICS1* gene expression and SA induction in Arabidopsis immune responses (Cui et al. 2017). EDS1 is the key signal protein in the interaction with Toll/interleukin 1 receptor (TIR)-nucleotide-binding/leucine-rich repeat (NLR) (TNL) receptors, during effector triggered immunity (ETI) for defense responses and cell death (Wiermer et al. 2005). Together with PAD4 or SENESCENCE-ASSOCIATED GENE101 (SAG101) it acts as a heterodimer to transcriptionally induce defense responses and ICS1-dependent SA synthesis (Cui et al. 2017; Rietz et al. 2011). The lesion phenotype of *fah1 fah2 loh2* was eliminated in *fah1 fah2 loh2 sid2-2* and *fah1 fah2 loh2 eds1-2* mutants, but still present in *fah1 fah2 loh2 pad4-1* though at a less severe level (Fig. 3), showing that the phenotype is dependent on SA and EDS1 but less on PAD4. Lapin and colleagues showed that EDS1 heterodimerizes with SAG101 and N REQUIRED GENE1 (NRG1) to promote TNL-dependent cell death in Arabidopsis, and that this strong cell death activity does not occur when PAD4 substitutes for SAG101 (Lapin et al. 2019). EDS1 and PAD4 instead mediate transcriptional promotion of defense reactions important for limiting bacterial growth. The distinction between both pathways seems not to be very strict and a small proportion of cell death can account from EDS1-PAD4 signaling. This could explain why no cell death is visible in *fah1 fah2 loh2 eds1-2* mutants, but is still found in *fah1 fah2 loh2 pad4-1* mutants.

Interestingly, not only the phenotype of the plants changed due to SA mutations but also the sphingolipid profile (Fig. 5). In *fah1 fah2 loh2 eds1-2*, *fah1 fah2 loh2 pad4-1* and especially in *fah1 fah2 loh2 sid2-2* plants the accumulation of Cers and LCBs is significantly reduced, showing that SA level has an influence on the sphingolipidome of Arabidopsis. Indeed, Shi et al. 2015 showed that SA influences sphingolipid fluxes in Arabidopsis seedlings by *in silico* flux balance analysis and suggested SA could also act upstream of LCB or Cer signals (Shi et al. 2015). Our data confirm now that SA levels in the plant influence LCB and Cer levels and that there might be a feedback loop between SA signaling and sphingolipid biosynthetic pathways.

Together our data strongly suggest that lesion formation as a result of changes in sphingolipid metabolism in *fah1 fah2 loh2* plants is SA-dependent and transduced via EDS1 signaling. SA signaling itself influences the accumulation of Cers and LCBs in Arabidopsis, suggesting a positive feedback loop between both pathways.

## Material and Methods

### Plant Material

The *Arabidopsis thaliana* plants used in this study were of the Columbia (Col-0) ecotype background. The seeds of the T-DNA insertion mutants: *fah1* (SALK_140660), *fah2* (SAIL_862_H01), *loh1* (SALK_069253), *loh2* (SALK_018608), *loh3* (SALK_150849), were obtained from the Nottingham Arabidopsis Stock Centre (NASC). All mutants are knock out mutants except *fah1* which is a knock down mutant (König et al. 2012; Ternes et al. 2011). The *sid2-2*, *eds1-2*, *pad4-1* mutants were kindly provided by Prof. Christiane Gatz (University of Goettingen, Germany). Double, triple and quadruple mutants were generated by crossing the above mutants. The homozygous insertion of T-DNA was verified by PCR analysis performed on genomic DNA. Primer sequences used for genotyping are provided in Table S1. For soil-grown plants, Arabidopsis seeds were sown on soil and then stratified at 4 °C for 2 days. Plants were grown at 22 °C under long day conditions (16 h light: 8 h dark) with a light intensity of 120-150 μmol m^-2^ s^-1^.

### Trypan Blue Staining

To visualize the cell death phenotype trypan blue staining was performed as described previously (Fernández-Bautista et al. 2016) and analyzed using a stereomicroscope Olympus SZX12 (Olympus, Hamburg, Germany)

### Quantification of SA and SAG

For absolute quantification of SA and SAG, 100 mg of frozen homogenized rosette leaf tissue were extracted with methyl-*tert*-butyl ether (MTBE) together with 10 ng D4-SA (C/D/N Isotopes Inc., Pointe-Claire, Canada), reversed phase-separated using an ACQUITY UPLC^®^ system (Waters Corp., Milford, MA, USA) and analysed by nanoelectrospray ionization (nanoESI) (TriVersa Nanomate^®^; Advion BioSciences, Ithaca, NY, USA) coupled with an AB Sciex 4000 QTRAP^®^ tandem mass spectrometer (AB Sciex, Framingham, MA, USA) employed in scheduled multiple reaction monitoring mode (Herrfurth and Feussner 2020). The reversed phase separation of SA and SAG was achieved by UPLC using an ACQUITY UPLC^®^ HSS T3 column (100 mm x 1 mm, 1.8 μm; Waters Corp., Milford, MA, USA). Solvent A and B were water and acetonitrile/water (90:10, *v/v*), respectively, both containing 0.3 mmol/l NH_4_HCOO (adjusted to pH 3.5 with formic acid). Mass transitions and optimized parameters for the detection of SA and SAG were as follows: 137/93 [declustering potential (DP) 25 V, entrance potential (EP) 6 V, collision energy (CE) 20 V] for SA, 141/97 (DP 25 V, EP 6 V, CE 22 V) for D_4_-SA and 299/137 (DP 30 V, EP 4 V, CE 18 V) for SAG. Quantification was carried out using a calibration curve of intensity (*m/z*) ratios of [unlabeled]/[deuterium-labeled] *vs*. molar amounts of unlabeled (0.3-1000 pmol). Data can be found in Supplemental Dataset II.

### Sphingolipid Analysis

The analysis of LCBs, LCB-Ps, Cers, GlucCers and GIPCs was performed according to a method previously described (Zienkiewicz et al. 2020). For extraction of sphingolipids,100 mg of frozen homogenized rosette leaf tissue were resuspended in propan-2-ol/hexane/water (60:26:16, *v*/*v*/*v*) and incubated at 60 °C for 30 min (Grillitsch et al. 2014). After centrifugation at 635 x g for 20 min, the supernatant was dried under a nitrogen stream and dissolved in tetrahydrofuran/methanol/water (4:4:1, *v*/*v*/*v*). For analysis of LCB-Ps, the resulting extract was measured after acetyl derivatization (Yanagawa et al. 2017). Sphingolipids were analyzed and resulting analytical data were processed as previously described with some modifications (Tarazona et al. 2015). The lipid extracts were reversed phase-separated using an ACQUITY UPLC^®^ system (Waters Corp., Milford, MA, USA) and analyzed by nanoESI (TriVersa Nanomate^®^; Advion BioSciences, Ithaca, NY, USA) coupled with an AB Sciex 6500 QTRAP^®^ tandem mass spectrometer (AB Sciex, Framingham, MA, USA) employed in scheduled multiple reaction monitoring mode. The reversed phase separation was achieved by UPLC using an ACQUITY UPLC^®^ HSS T3 column (100 mm x 1 mm, 1.8 μm; Waters Corp., Milford, MA, USA). Solvent A and B were methanol/20 mM ammonium acetate (3:7, *v/v*) and tetrahydrofuran/methanol/20 mM ammonium acetate (6:3:1, *v/v/v*), respectively, both containing 0.1 %, *v/v* acetic acid. Data can be found in Supplemental Dataset I.

## Acknowledgements

We are grateful to Dr. Tegan Haslam for critical reading the manuscript and to Sabine Freitag for her expert technical assistance.

