## Supplemental Files Fig S1-S3 Tab S1 for "Sphingolipid-Induced Programmed Cell Death Is a Salicylic Acid and EDS1-Dependent Phenotype in Arabidopsis"

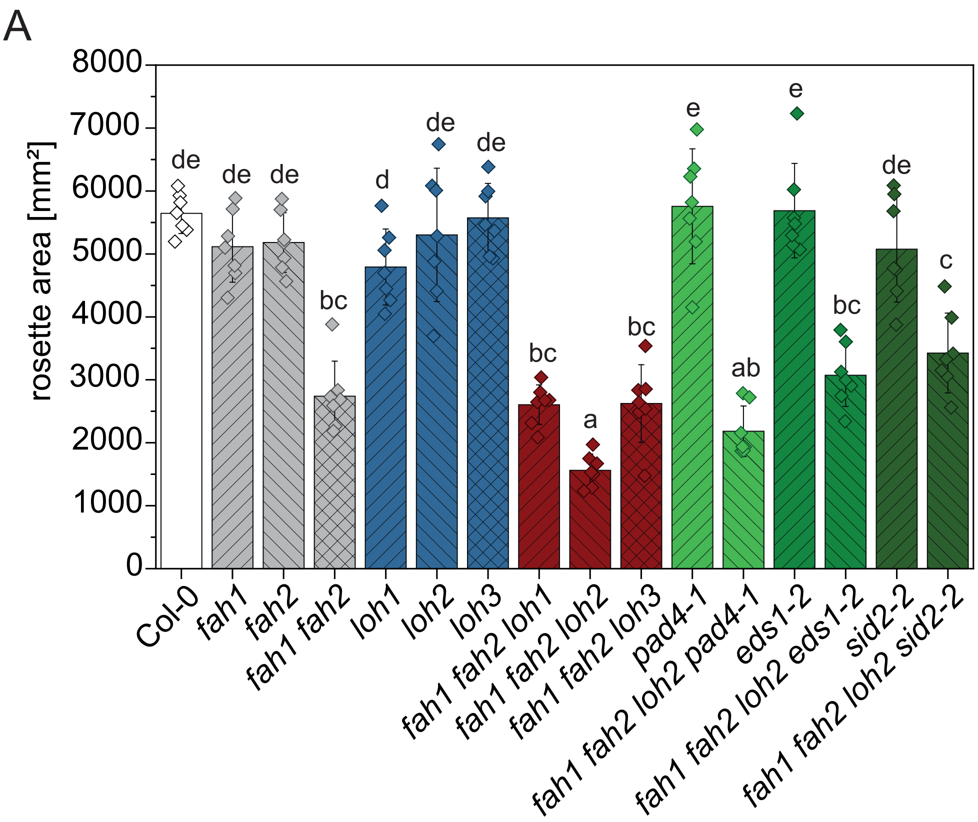


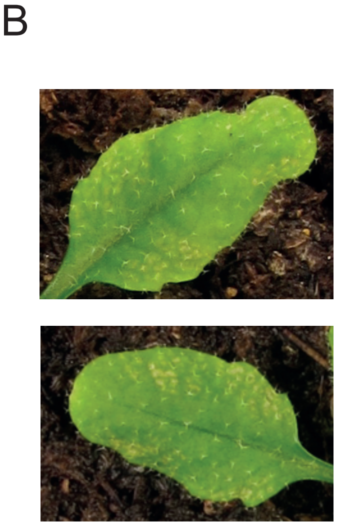


**Fig. S1** (A) Crossing SA synthesis and signaling mutants with plants defective in *Fatty Acid Hydroxylase1* and *2* and CerS *LOH2* partially complements rosette area reduction. Rosette area was quantified from 35-day-old plants grown under long day conditions. Pictures of each plant were taken from above and the projected rosette area was determined by a software program provided by datInf (Tübingen). Values represent the mean ±SD of 13 biological replicates out of two independent experiments. Statistical analysis was performed with log-transformed data by one-way analysis of variance (ANOVA) with Tukey's *post hoc* test ($P$<0.05). Different letters indicate significant differences with $P$<0.05. (B) Visible lesions in *fah1 fah2 loh2* mutant plants.

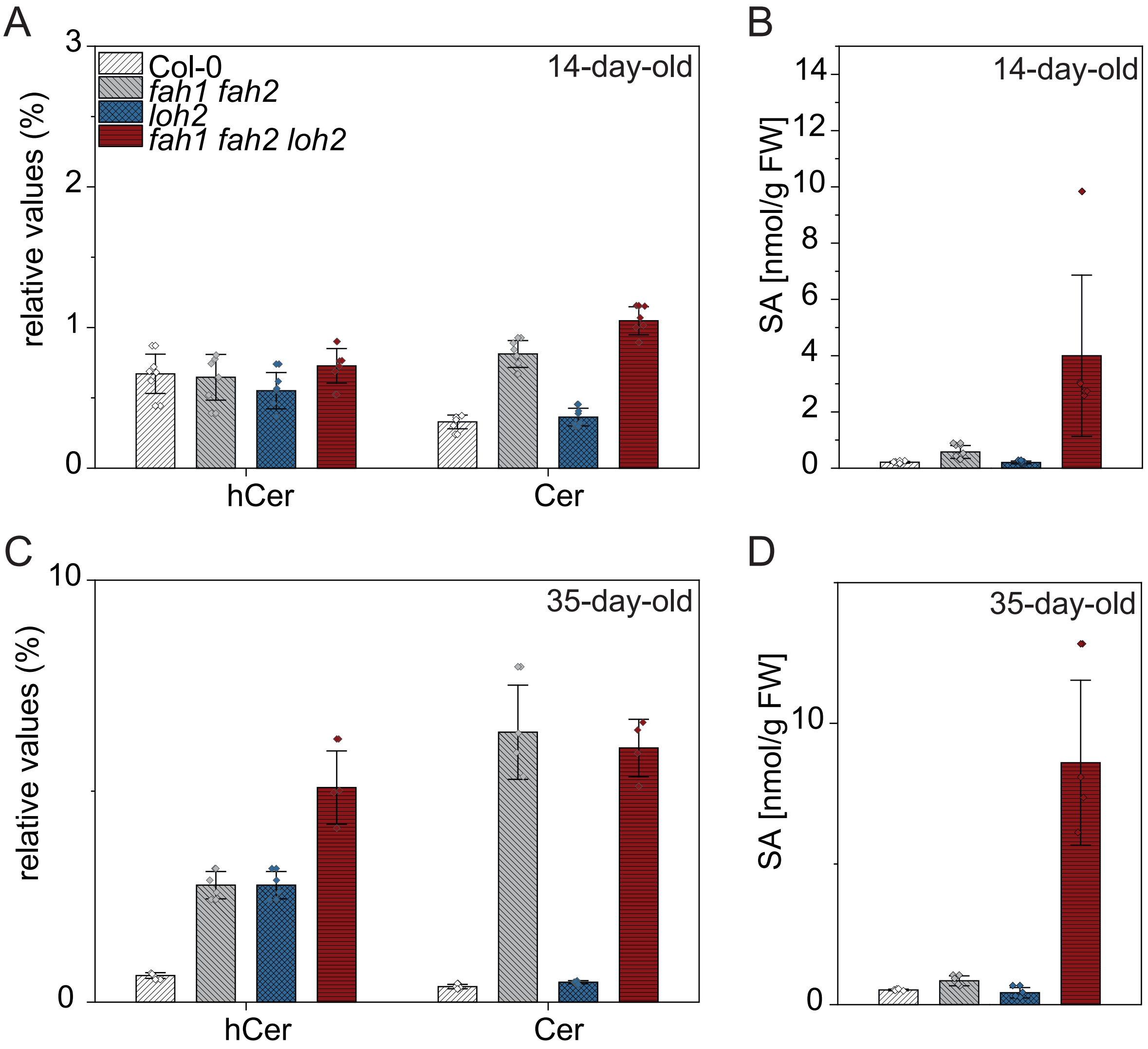


**Fig. S2** SA (B, D), Cer and hCer (A,C) content in 14- and 35-day-old plants. Rosette leaves of plants grown under long day conditions were extracted and analyzed. Values represent the mean ±SD of six (14-day-old) or four (35-day-old) biological replicates (n=6 and n=4). The experiment was repeated once with similar tendencies.

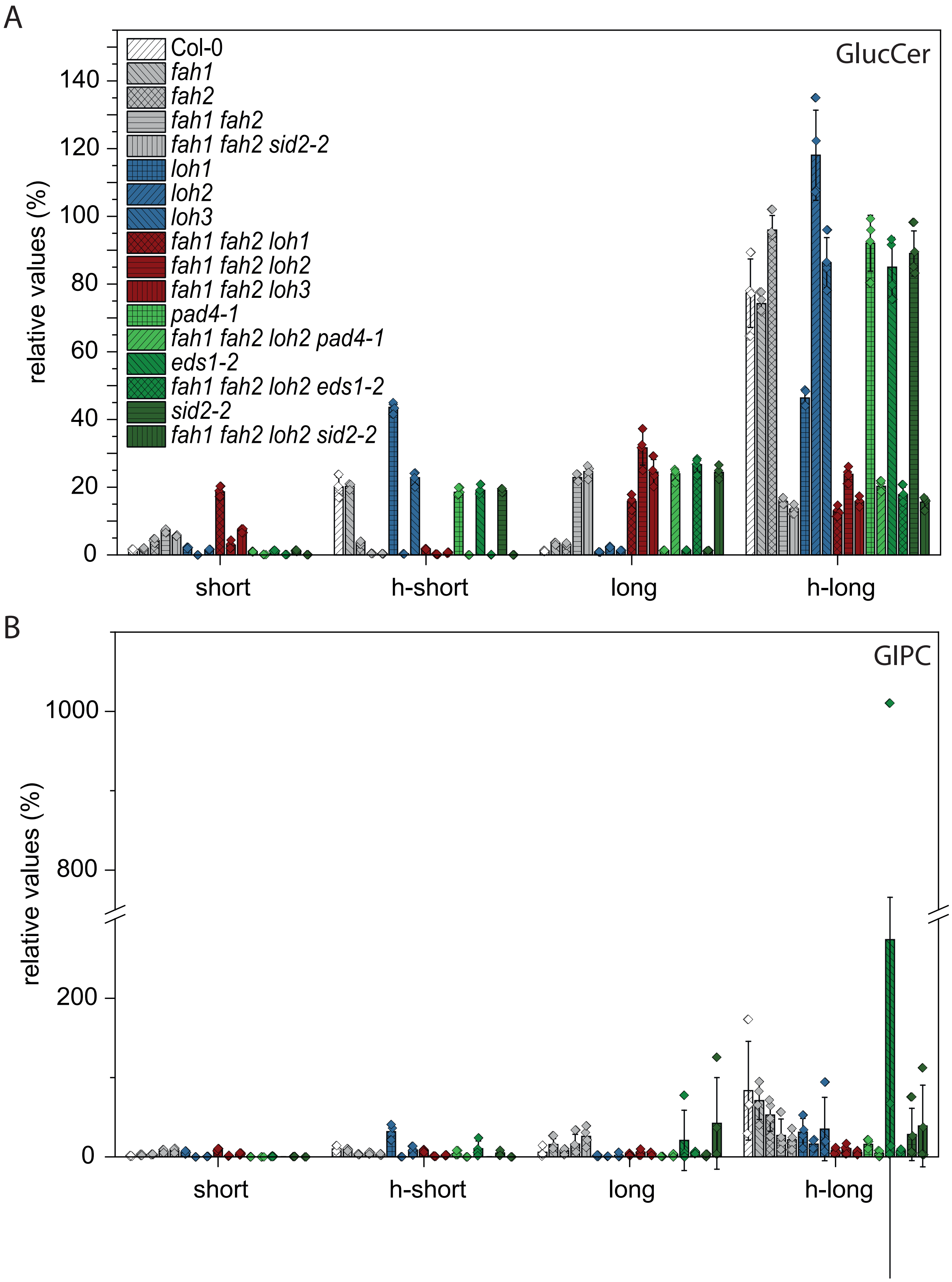


**Fig. S3** GlucCer (A) and GIPC (B) profiles in crosses of *fah1 fah2* double and *fah1 fah2 loh2* triple mutants with SA synthesis (*sid2-2*) and signaling mutants (*eds1-2, pad4-1*). Rosette leaves of 35-day-old plants grown under long day conditions were extracted and analyzed. LCFA, Cers with C16-C18 FA moiety; hLCFA, Cers with C16-C18 α-hydroxylated FA moiety; VLCFA, Cers with C20-C28 FA moiety; hVLCFA, Cers with C20-C28 α-hydroxylated FA moiety Values represent the mean ±SD of four biological replicates (n=4). The experiment was repeated once with similar tendencies.

**Table S1:** Gene specific primers (LP – left primer, RP- right primer) and T-DNA left border primers (LBb1.3 and LB3) used for genotyping of plants.

| Primer Name | Primer Sequence (5`→3`) | Note | Mutant Line |
|---|---|---|---|
| *FAH1* LP | ACATGTTTTCGAGACACCTTCC | T-DNA insertion | SALK_140660 |
| *FAH1* RP | CACAATCAACGGTCCAGATTCC | | |
| *FAH2* LP | CATCCAAACCAAGAGTTACTGG | T-DNA insertion | SAIL_862_H01 |
| *FAH2* RP | GTCTAGAGATGGTTTTCCTTACC | | |
| *LOH1* LP | TTCACCTGCTTTATTGATGGG | T-DNA insertion | SALK_069253 |
| *LOH1* RP | TTGATCAGGCCAAATCTGATC | | |
| *LOH2* LP | CCAGGAGTTCAATGCTTCAAC | T-DNA insertion | SALK_018608 |
| *LOH2* RP | ATGCTGCTTGTGACTTTTTCG | | |
| *LOH3* LP | TTCTCGATTTTGTGCTTGCTC | T-DNA insertion | SALK_150849 |
| *LOH3* RP | CACCCATTAGCGATCAGAATC | | |
| *PAD4* p1 | GCGATGCATCAGAAGAG | digested with Faq1 | *pad4-1* |
| *PAD4* p2 | TTAGCCCAAAAGCAAGTATC | | |
| 105/E2 | ACACAAGGGTGATGCGAGACA | Deletion | *eds1-2* |
| *EDS* 4 | GGCTTGTATTCATCTTCTATC C | | |
| *EDS* 6 | GTGGAAACCAAATTTGACATTAG | | |
| *SID2* LP | CAAGAGACAACATTGCTTTC | Deletion | *sid2-2* |
| *SID2* RP | CAGAAGGTTTTTGTCTTCAACG | | |
| LBb1.3 | ATTTTGCCGATTTCGGAAC | | For SALK Line |
| LB1 | GCCTTTTCAGAAATGGATAAATAG CCTTGCTTCC | | For SAIL Line |
